# Reversal of cardiac and skeletal manifestations of Duchenne muscular dystrophy by cardiosphere-derived cells and their exosomes in *mdx* dystrophic mice and in human Duchenne cardiomyocytes

**DOI:** 10.1101/128900

**Authors:** Mark A. Aminzadeh, Russell G. Rogers, Kenneth Gouin, Mario Fournier, Rachel E. Tobin, Xuan Guan, Martin K. Childers, Allen M. Andres, David J. Taylor, Ahmed Ibrahim, Xiangming Ding, Angelo Torrente, Joshua M. Goldhaber, Ronald A. Victor, Roberta A. Gottlieb, Michael Lewis, Eduardo Marbán

**Affiliations:** Cedars-Sinai Heart Institute, Los Angeles, CA USA; Institute for Stem Cell and Regenerative Medicine, University of Washington, Seattle, WA USA; UCLA Technology Center for Genomics & Bioinformatics, Los Angeles, CA USA

## Abstract

Genetic deficiency of dystrophin leads to disability and premature death in Duchenne muscular dystrophy, affecting the heart as well as skeletal muscle. Here we report that cardiosphere-derived cells (CDCs), which are being tested clinically for the treatment of Duchenne cardiomyopathy, improve cardiac and skeletal myopathy in the *mdx* mouse model of DMD and in human Duchenne cardiomyocytes. Injection of CDCs into the hearts of *mdx* mice augments cardiac function, ambulatory capacity and survival. Exosomes secreted by human CDCs reproduce the benefits of CDCs in *mdx* mice and in human Duchenne cardiomyocytes. The findings further motivate the testing of CDCs in Duchenne patients, while identifying exosomes as next-generation therapeutic candidates.

## INTRODUCTION

Absence of dystrophin in Duchenne muscular dystrophy (DMD) leads to membrane fragility and secondary damage to muscle (both skeletal and cardiac)(1). Early disability is due predominantly to the skeletal myopathy, but heart failure is the most common cause of death(2). Cardiosphere-derived cells (CDCs) may represent a viable therapeutic option(3-5). CDCs are progenitor cells intrinsic to the heart; in clinical trials after myocardial infarction, CDCs promote cardiomyogenesis and reverse established scar (6, 7). Multiple lines of evidence now indicate that most of the beneficial effects of CDCs are indirect. In the extreme, allogeneic CDCs are cleared completely within several weeks, but their functional and structural benefits persist at least 6 months (8). CDCs secrete diffusible factors that promote angiogenesis, recruit endogenous progenitor cells, and coax surviving heart cells to proliferate (9, 10); transplanted CDCs suppress maladaptive remodeling (11), apoptosis (10, 12), fibrosis (13), and inflammation after myocardial infarction (13) and in nonischemic cardiomyopathy (14). These diverse mechanisms appear to be mediated via the secretion of exosomes laden with noncoding RNA including microRNAs (miRs) (15), consistent with the notion that exosomes contain a plethora of bioactive molecules which target multiple signaling pathways synergistically (16). In a murine model of myocardial infarction, CDC-secreted exosomes (CDC-exosomes) mimic the functional and structural benefits of CDCs, while blockade of exosome biosynthesis renders CDCs ineffective (15). Given the clinical data with CDCs, and the complementarity between their therapeutic actions and the pathophysiological processes underlying Duchenne cardiomyopathy (oxidative stress (17, 18), inflammation (19), fibrosis (20), and mitochondrial dysfunction (21)), we reasoned that CDCs and their exosomes might be useful in treating Duchenne cardiomyopathy. Our early work reported in abstract form (22, 23) revealed striking phenotypic correction by CDCs in *mdx* dystrophic mice, motivating the ongoing HOPE-Duchenne clinical trial (24) of CDCs in DMD patients. Initially, we had not aspired to restore skeletal muscle function, but merely to offset the pathophysiological consequences of dystrophin deletion in the heart. We now report that CDCs and their secreted exosomes potently improve skeletal muscle structure and function, contributing to major systemic benefits after injection of CDCs into the heart.

## RESULTS

### CDC transplantation in *mdx* hearts

Intramyocardial injection of first and second (lower) doses of CDCs into the hearts of *mdx* mice improved left ventricular function (as manifested by ejection fraction [EF]) and volumes, relative to placebo, for at least 6 months (Fig. 1A and fig. S1A). The CDC-induced improvement in EF persisted beyond the point at which no surviving CDCs were detectable in *mdx* hearts (3 weeks after CDC delivery; fig. S1B). In addition to improving EF, CDC injection enhanced ambulatory function (Fig. 1B). Ten-month-old wild-type mice (WT) and *mdx* mice (distinct from the *mdx* mice studied in Fig. 1A) were subjected to weekly high-intensity treadmill exercise, starting 3 weeks after single-dose CDC or vehicle administration. CDC-treated *mdx* mice showed a substantial increase in maximal exercise capacity, relative to vehicle-treated *mdx* mice, over the 3 mos that exercise capacity was measured; survival also differed in the two groups (Fig. 1C). By ∼23 mos of age, all vehicle-treated *mdx* mice had died, whereas > 50% of CDC-treated *mdx* mice remained alive (Fig. 1C). In investigating mechanism, we first studied known (anti-oxidative, anti-inflammatory, anti-fibrotic, and cardiomyogenic) effects of CDCs (3-15). Injection of CDCs led to major changes in the expression of genes related to oxidative stress, inflammation and mitochondrial integrity (fig. S2). The Nrf2 anti-oxidant pathway was activated in CDC-treated *mdx* heart (Fig. 1D). Nrf2 is normally repressed by Keap1, but oxidative stress (as well as Nrf2 phosphorylation by protein kinases such as Akt) causes dissociation of the Nrf2-Keap1 complex, culminating in nuclear translocation of Nrf2 and transcriptional activation of antioxidant enzymes (25). In *mdx* hearts, levels of phosphorylated Akt (Fig. 1E), total Nrf2 (Fig. 1F) and nuclear Nrf2 (Fig. 1G) were high (as expected in response to oxidative stress); CDC treatment further increased their protein levels (Figs. 1D-G) and those of downstream gene products (heme oxygenase-1 [HO-1], catalase, superoxide dismutase-2 [SOD-2], and the catalytic subunit of glutamate-cysteine ligase [GCLC]; Fig. 1H & fig. S2E). Concomitantly, oxidative stress was attenuated, as evidenced by a profound reduction of malondialdehyde adducts (Fig. 1I). Histological analysis revealed extensive fibrosis in vehicle-treated *mdx* hearts, but much less in CDC-treated *mdx* hearts (comparable to an age-matched wild-type [WT] control; fig. S3A). Likewise, CDC treatment largely reversed the accumulation of collagens I and III in *mdx* heart tissue 3 weeks after treatment (fig. S3B). CDCs inhibited the inflammation (Figs. 1J,K) and mitochondrial dysfunction (Figs. 1L-N) characteristic of *mdx* cardiomyopathy. NFκB, the master regulator of pro-inflammatory cytokines and chemokines (26), was activated in vehicle *mdx* hearts (Fig. 1K, top panel). Increases in phosphorylated IκB and nuclear p65 were accompanied by upregulation of MCP1 (monocyte chemoattractant protein1) and accumulation of CD68^+^ macrophages and CD3^+^ T cells (Fig. 1K, bottom panel). CDC treatment reversed activation of NFκB and decreased the number of inflammatory cells in *mdx* hearts 3 weeks after CDC injection (Figs. 1J,K; figs. S4, S5). Mitochondrial structure and function are abnormal in muscular dystrophy-associated heart failure (21). Whole-transcriptome analysis revealed major changes in the expression of genes related to mitochondrial integrity in *mdx* hearts (fig. S2B). Consistent with this finding, CDCs restored mitochondrial ultrastructure (Fig. 1L), increased mitochondrial DNA copy numbers (but not mitochondrial number; fig. S6), augmented levels of respiratory chain subunits (Fig. 1M) and normalized the deficient respiratory capacity of isolated *mdx* mitochondria (Fig. 1N). Of note, the salutary mitochondrial changes were associated with upregulation of antioxidant enzymes and reductions of oxidative stress and inflammation (Figs. 1D-k; figs. S2C-E, S5). We also probed the effects of CDCs on cardiomyogenesis. Vehicle-treated *mdx* hearts exhibited a modest increase in the numbers of cycling (Ki67^+^) and proliferating (aurora B^+^) cardiomyocytes (fig. S7), presumably as a compensation for ongoing cardiomyocyte loss. CDCs are known to increase endogenous cardiomyogenesis in ischemic (7, 11, 12, 27, 28) and non-ischemic models (14). Similar effects were seen in the *mdx* heart: CDC treatment promoted cardiomyocyte cycling and proliferation, as evidenced by a marked increase in Ki67^+^ and aurora B^+^ cardiomyocytes (fig. S7).

**Figure 1.**
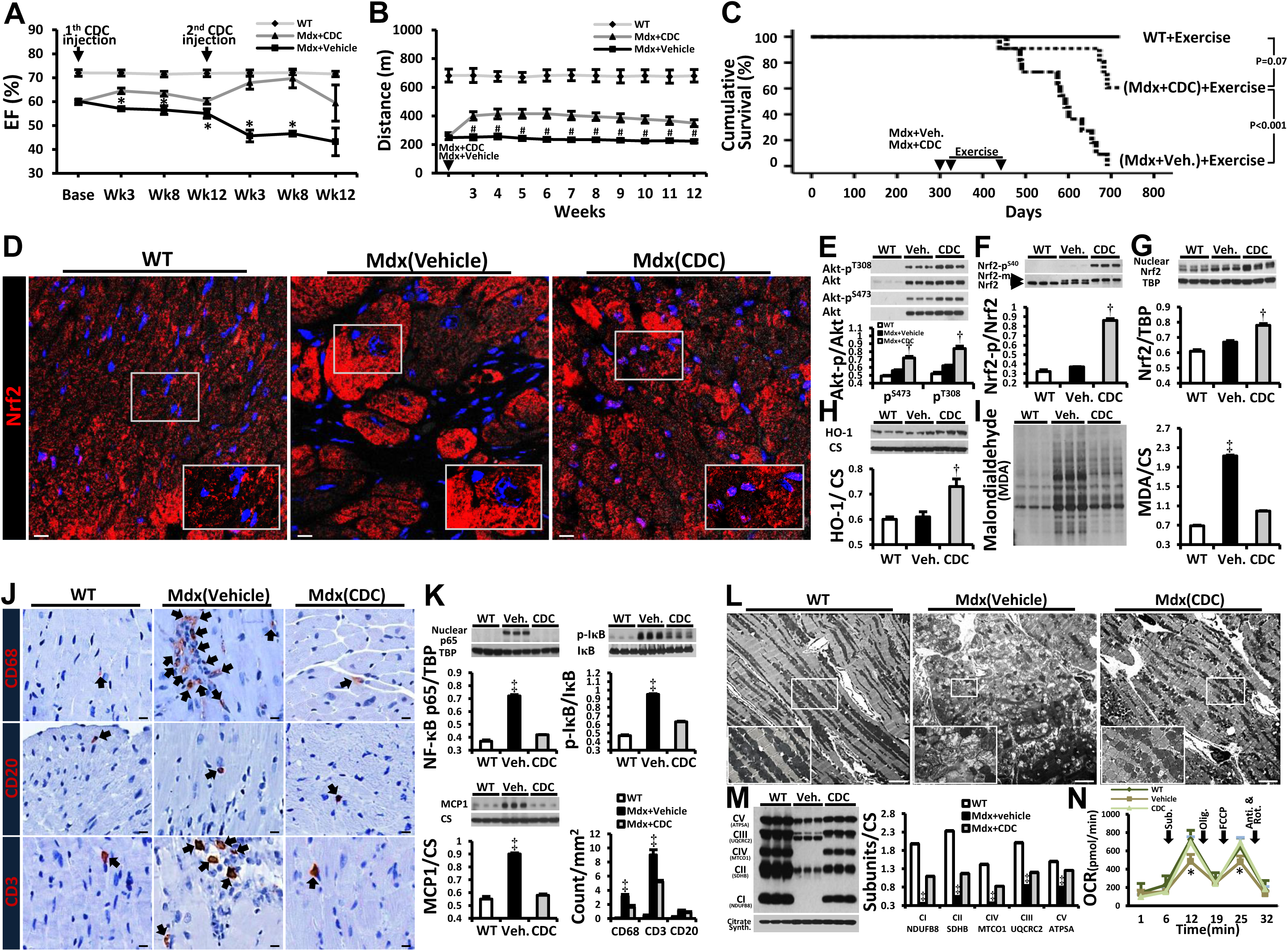
CDC transplantation into *mdx* hearts. Function, survival, antioxidant pathways, inflammation and mitochondrial dysfunction improved by CDC transplantation into *mdx* mice. A: Ejection fraction (EF) in CDC-injected *mdx* mice (Mdx+CDC) and vehicle-injected *mdx* mice (Mdx+Vehicle) in response to injections at baseline (10 mos of age) and 3 months later (WT: n=7; Mdx+Vehicle & Mdx+CDC: n=12 each). B: Exercise capacity in mice subjected to weekly high-intensity treadmill exercise, starting 3 weeks after single-dose CDC or vehicle administration (WT: n=7; Mdx+Vehicle & Mdx+CDC: n=11 each). Cardiac (A) and treadmill (B) experiments were performed separately on different groups of experimental mice. C: Kaplan-Meier analysis of survival in the same animals as B shows lower survival in vehicle-treated *mdx* mice than in CDC-treated *mdx* mice or wild-type controls (p< 0.001, log rank test); the latter two groups, however, were statistically comparable. D: Immunohistochemical images of Nrf2 in *mdx* mouse hearts 3 weeks after administration of vehicle or CDCs. Age-matched wild-type mice (WT) served as control. E-I: Western blots and pooled data for protein abundance of phospho-Akt [Akt-p^T308^, Akt-p^S473^, (E)], cytoplasmic phospho-Nrf2 [Nrf2-p^S40^, (F)], nuclear Nrf2 (G), Nrf2 downstream gene product, heme oxygenase-1 [HO-1; (H)], and malondialdehyde protein adducts(I) in *mdx* mouse hearts 3 weeks after administration of vehicle or CDCs (WT: n=4; Mdx+Vehicle & Mdx+CDC: n=6 each). J: Immunohistochemical images of hearts stained for inflammatory cell markers CD68, CD20 and CD3; K: Western blots, pooled data and bar graph (lower right) representing protein abundance of nuclear p65, p-IκB (NF-κB pathway) and MCP1 (Monocyte Chemoattractant Protein1) and average number of indicated inflammatory cells and in *mdx* mouse hearts. L: Transmission electron microscopy (TEM) images from *mdx* mouse hearts 3 weeks after administration of vehicle (Mdx+Vehicle) or CDCs (Mdx+CDC). Age-matched wild-type mice (WT) served as control. M&N: Representative western blots and pooled data for mitochondrial respiratory chain subunits in WT and vehicle/CDC *mdx* heart tissues (M) and oxygen consumption rate (OCR) of mitochondria isolated from the hearts of WT and CDC- or vehicle-treated *mdx* mice (N) 3 weeks after treatment (WT: n=3; Mdx+Vehicle & Mdx+CDC: n=8 each). Substrates (pyruvate, malate and ADP), a selective uncoupler (FCCP) and blockers (Oligomycin [Olig.]; Antimycin & Rotenone [Anti. & Rot.]) of oxidative phosphorylation were applied when indicated. Pooled data are means ± SEM. CM: Cardiomyocyte; CMs: Cardiomyocytes.*\*P< 0.05 vs. Mdx+CDC; #P< 0.005 vs. Mdx+CDC; †P< 0.05 vs. Mdx+Vehicle and WT (wild type mice); ‡P< 0.002 vs. Mdx+CDC and WT (wild type mice). Scale bars: 10μm (D,J); 5 μm (L).*

### CDC-exosome transplantation in *mdx* hearts

CDC-exosomes mimic the functional and structural benefits of CDCs in a murine model of myocardial infarction (15). In *mdx* mice, likewise, exosomes, isolated from media conditioned by hypoxic CDCs, reproduced the benefits of CDCs (Figs. 2A-C, fig. S8A-D). Two repeat doses of human CDC-exosomes (separated by 3 months) led to sustained improvement in EF, relative to vehicle injection (Fig. 2A & fig. S8B), with a minimal but detectable humoral response in the non-immunosuppressed *mdx* mice (fig. S8C). Collagen I and III levels decreased (Fig. 2B) while cycling (Ki67^+^, Fig. 2C upper row) and proliferating (aurora B^+^, Fig. 2C lower row) cardiomyocytes increased in CDC-exosome-injected *mdx* hearts. The effects of CDC-exosomes were mediated at least in part via clathrin-mediated endocytosis by the surrounding myocardium (fig. S8D).

**Figure 2.**
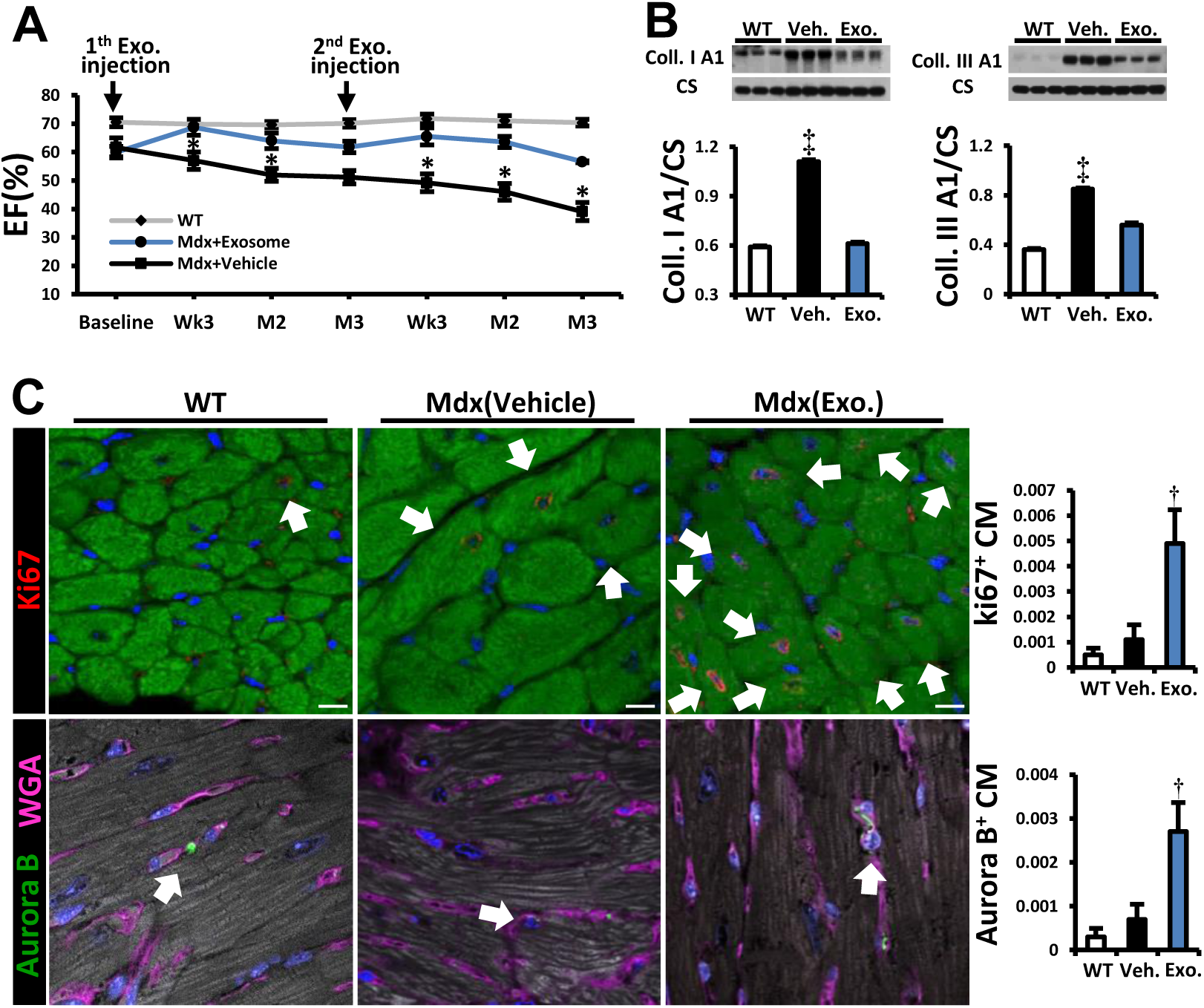
CDC-exosome injection into *mdx* hearts reproduces benefits of CDCs. A: Sustained functional benefit for at least 3 months with each of two sequential CDC exosome injections in *mdx* mice (n=11). B: Western blots and pooled data for cardiac collagen IA and IIIA. C: Immunohistochemical images and pooled data (WT [wild type; n=4], vehicle and CDC-exosome-treated [Mdx (XO)]; n=6 each) from *mdx* mouse hearts stained for Ki67 [upper row] and Aurora B [lower row]). Arrows point to Ki67^+^ (upper row) and Aurora B^+^ (lower row) cardiomyocytes. Data are means ± SEM; *\*P< 0.05 vs. Mdx+exosome; ‡P< 0.01 vs. Mdx+exosome and WT (wild type mice); †P< 0.02 vs. Mdx+Vehicle and WT (wild type mice); scale bar: 10μm.*

### Remote effects of CDC transplantation in *mdx* heart

Intramyocardial injection of CDCs and their exosomes improved Duchenne cardiomyopathy, reversing key pathophysiological processes in the *mdx* mouse heart (Figs. 1, 2). These changes were associated with a substantial increase in exercise capacity, which was disproportionate to the improvement in cardiac function: EF increased by < 10% (Fig. 1A), while ambulatory capacity doubled (Fig. 1B). To further evaluate the mechanism of enhanced exercise capacity in CDC-treated *mdx* mice, we isolated and examined three distinct skeletal muscles: the diaphragm (DIA, a key respiratory muscle), and two limb muscles (soleus and extensor digitorum longus [EDL], representative of slow and fast twitch muscles, respectively) 3 weeks after intramyocardial injection of CDCs or vehicle. Whole-transcriptome analysis in DIA revealed down-regulation of pathways related to intracellular [Ca^2+^] excess, oxidative stress and inflammation after intramyocardial CDC injection (Fig. 3A & fig. S9). Decreases in malondialdehyde protein adducts (Fig. 3B), repressed NFκB, reduced infiltration of inflammatory cells (Fig. 3C) and diminished fibrosis (Fig. 3D) paralleled a marked improvement in the contractile function of DIA (Figs. 3E-G). Similarly, soleus (Figs. 3H-K) and EDL (figs. S10A) showed notable improvements at transcriptomic, histologic and functional levels; soleus fibrosis (fig. S10B) was attenuated and contractile force (Figs. 3I-K) was augmented. Changes in gene expression in DIA and soleus were significantly correlated (Fig. 3L).

**Figure 3.**
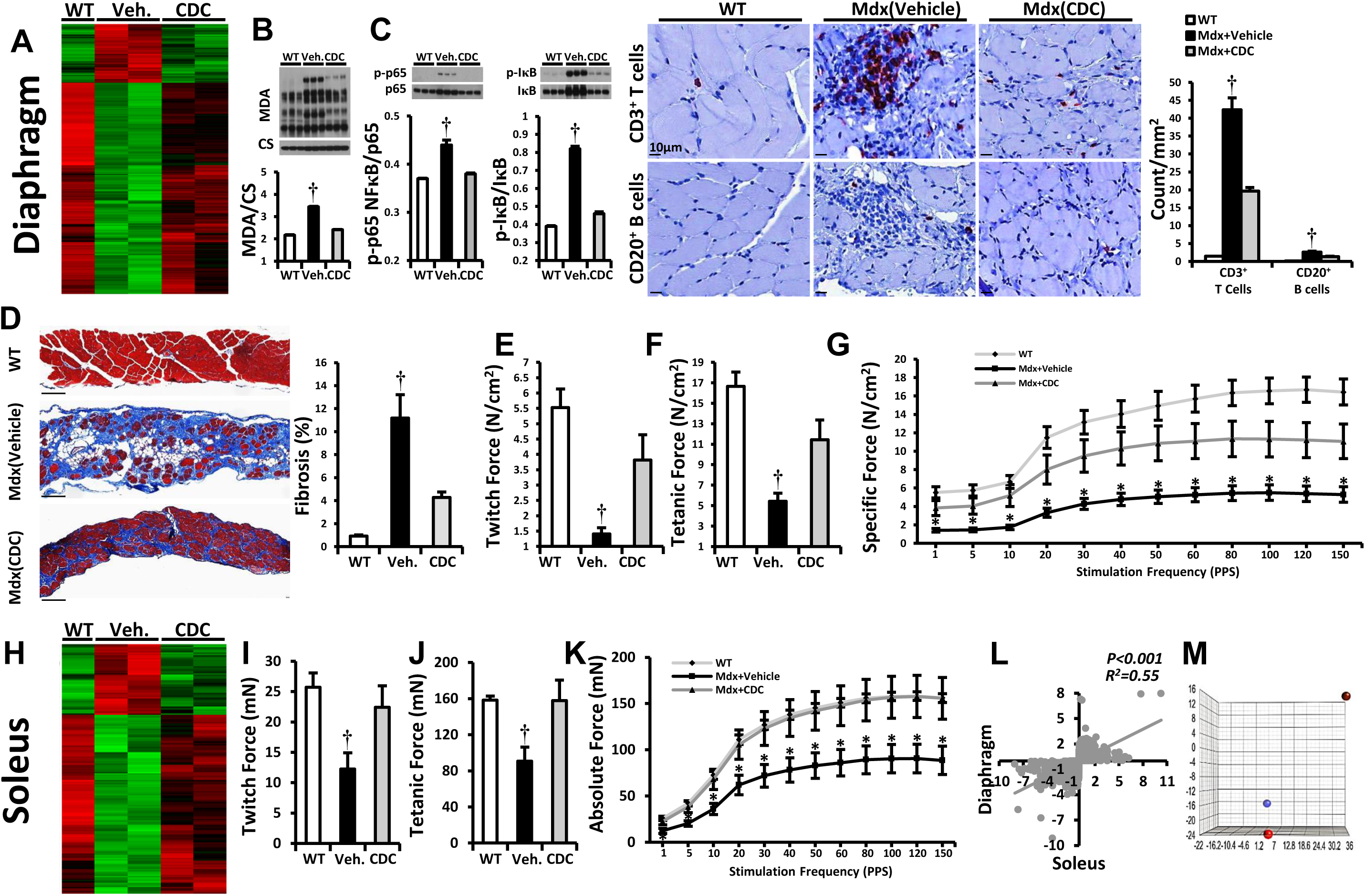
CDC transplantation in *mdx* hearts conferred beneficial effects on diaphragm and soleus muscles. A: 2-Dimensional hierarchical clustering using genes with at least 2 times fold change difference between vehicle/CDC *mdx* diaphragms. B&C: Western blots and pooled data for protein abundance of malondialdehyde protein adducts (B), cytoplasmic p-p65 and p-IκB (C; NF-κB pathway; WT: n=4; Vehicle & CDC: n=6 each) and immunohistochemical images of diaphragm stained for inflammatory cell markers CD20 and CD3; bar graph represents average number of indicated inflammatory cells 3 weeks after administration of vehicle or CDCs into *mdx* hearts. D: Representative Masson trichrome images and morphometric analysis in diaphragms 3 weeks after administration of vehicle or CDCs into the hearts of *mdx* mice. E-G: *in vitro* measurement of diaphragm contractile properties: twitch force (E), maximum tetanic force (F) and specific force (G) 3 weeks after CDC/vehicle *mdx* heart treatments. H: 2-Dimensional hierarchical clustering using genes with at least 2 times fold change difference between vehicle/CDC *mdx* soleus. I-K: *ex vivo* measurement of soleus contractile properties: twitch force (I), maximum tetanic force (J) and absolute force (K) 3 weeks after CDC/vehicle treatment of *mdx* hearts. L: Correlation of fold changes in expression of same genes in diaphragm and soleus 3 weeks after intramyocardial CDC injection in *mdx* mice. M: Three-dimensional plot depicting principal components analysis (PCA) of RNA-seq expression data from exosomes isolated from hypoxic conditioned media and effluents of CDC-or vehicle-treated *mdx* hearts. The effluent of isolated *mdx* hearts undergoing Langendorff perfusion was collected for exosomes isolation and subsequent RNA-seq 3 days after intramyocardial CDC/vehicle injection. PCA analysis showed clustering of CDC-exosomes (red) with exosomes isolated from effluent of CDC *mdx* hearts (blue), but not vehicle-injected *mdx* hearts (stippled), indicating that CDC-exosomes were shed from *mdx* hearts at least 3 days after intramyocardial CDC injection. Effluents of *mdx* hearts from same group were pooled (n=3 for each group). Data are means ± SEM; *\*P< 0.05 vs. Mdx+CDC; †P< 0.05 vs. Mdx+CDC and WT (wild type mice).*

As a basis for the remote effects of intramyocardial injection of CDCs on skeletal muscle, we considered the possibility that exosomes secreted by CDCs lodged in the heart might exit in the venous effluent and exert remote signaling. Principal components analysis revealed that CDC-exosomes were very similar in their RNA contents to the exosomes isolated from effluents of isolated CDC-treated *mdx* hearts, but quite distinct from exosomes in the effluents of vehicle-treated *mdx* hearts 3 days after intramyocardial CDC injection (Fig. 3M), pinpointing exosomes as likely mediators of the secondary systemic effects. Such secondary effects are extensive: whole transcriptome analysis of liver (fig. S11A) 3 weeks after intramyocardial CDC injection revealed downregulation of inflammatory pathways in liver analogous to what we found in heart and skeletal muscle. Thus, CDCs’ secondary effects are not restricted to muscle.

### Systemic CDC-exosome injection

To further evaluate the potential of exosomes to mediate systemic benefits, we injected CDC-exosomes into the left ventricular cavity of *mdx* hearts. Six hours post-injection, fluorescently-labeled CDC-exosomes were evident not only in the heart and skeletal muscle (Fig. 4A), but also in brain, liver, lung, spleen, gut and kidneys (fig. S11B). Changes in *mdx* heart (Figs. 4B-E), diaphragm (DIA; Figs. 4F-H) and soleus (Figs. 4I-K) 3 weeks after intraventricular CDC-exosome injection mimicked the modifications seen in these organs after intramyocardial CDC injection (Fig. 3). Taken together, the results in Figs. 1-4 implicate CDC-exosomes as mediators of the local and remote effects of intramyocardial CDC injection.

**Figure 4.**
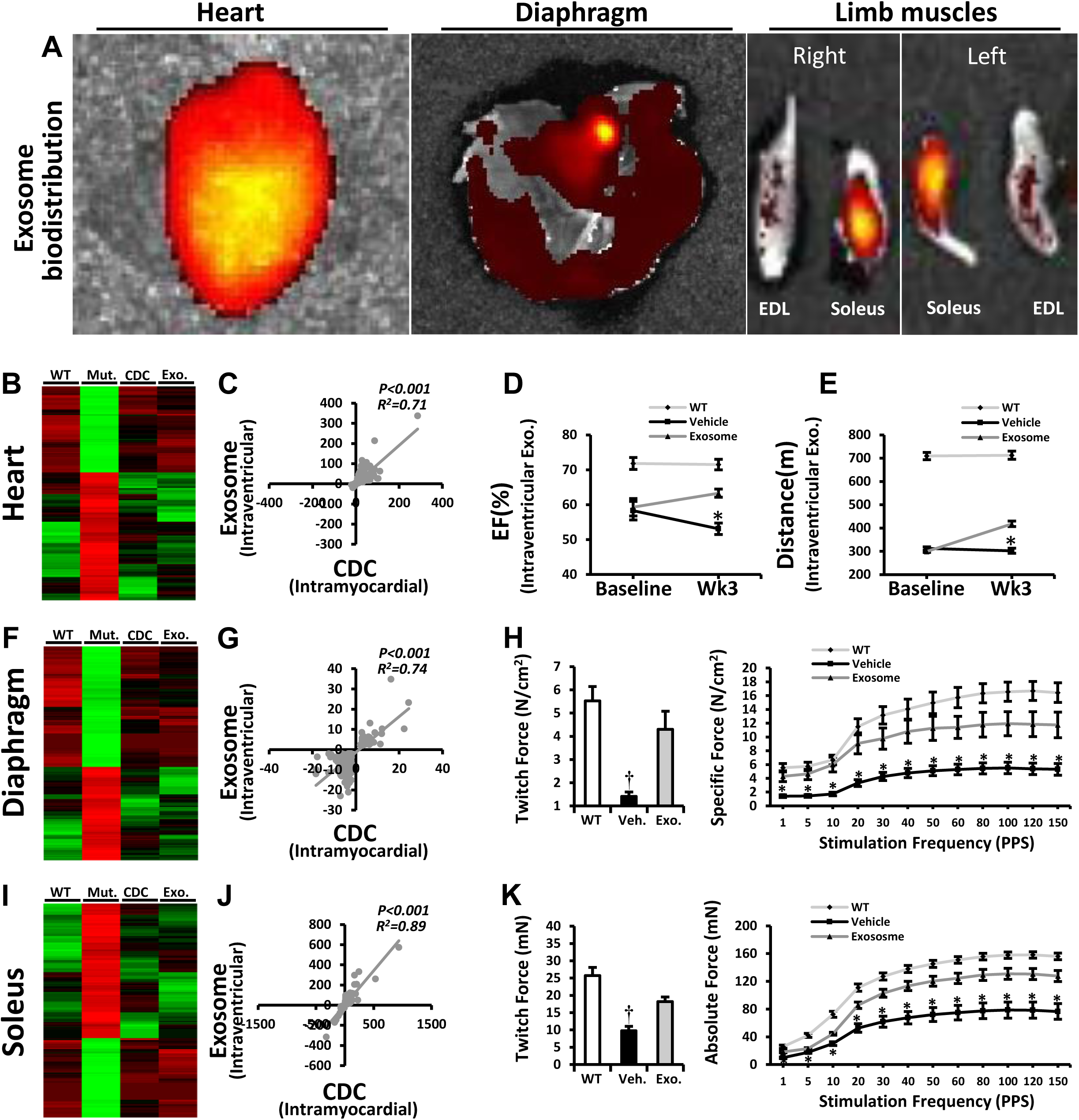
Systemic CDC-exosome injection mimicked the cardiac and the remote effects of intramyocardial CDC injection in *mdx* mice. A: Systemic biodistribution of CDC-exosomes after intraventricular injection in *mdx* mice. CDC-exosomes were stained with fluorescent lipid dye and tracked 6 hours later using bioluminescence imaging. B: 2-Dimensional hierarchical clustering using genes from hearts of non-treated *mdx* mice and of *mdx* mice treated intramyocardially with CDCs or intraventricularly with CDC-exosomes. Genes with at least 2-fold differences with corresponding transcripts in non-treated *mdx* mice were included. C: Correlation of fold changes in expression of same genes 3 weeks after intramyocardial CDC injection or intraventricular CDC-exosome injection in *mdx* hearts. D&E: EF and exercise capacity in *mdx* mice 3 weeks after intraventricular injection of vehicle/CDC-exosome (WT: n=5; Mdx+Vehicle & Mdx+CDC-exosome: n=9 each). F: 2-Dimensional hierarchical clustering using genes from diaphragm of non-treated *mdx* mice and of *mdx* mice treated intramyocardially with CDCs or intraventricularly with CDC-exosomes. Genes with at least 2-fold differences with corresponding genes in non-treated *mdx* mice were included. G: Correlation of fold changes in expression of the same genes in diaphragm 3 weeks after intramyocardial CDC injection or intraventricular CDC-exosomes injection. H: diaphragm contractile properties 3 weeks after intraventricular CDC-exosome injection. I-K: 2-Dimensional hierarchical clustering (I), correlation analysis (J) and contractile properties (K) from soleus muscle 3 weeks after intraventricular CDC-exosome injection. Data are means ± SEM; *\*P< 0.05 vs. Mdx+CDC-exosome; †P< 0.05 vs. Mdx+CDC-exosome and WT (wild type mice).*

### CDC-exosome injection into *mdx* skeletal muscle

To investigate primary effects on skeletal muscle, we injected CDC-exosomes directly into the soleus in *mdx* mice. Histological analysis revealed a paucity of surviving myofibers in vehicle-injected *mdx* soleus relative to wild-type controls, and those that remained were hypertrophic (Fig. 5A). CDC-exosomes markedly increased the total number of myofibers and shifted the size distribution to smaller diameters, indicative of myofiber proliferation 3 weeks after injection (Fig. 5B). Consistent with this interpretation, the number of MyoD^+^ cells was augmented after CDC-exosome injection (Figs. 5A,C), with increased tissue levels of MyoD and myogenin, the major transcription factors orchestrating myoblast determination and differentiation, respectively (29) (Fig. 5D). In physiological muscle growth, IGF-1 is commonly implicated as an upstream signal (30), but the effects of CDC-exosomes on *mdx* soleus muscle were independent of IGF-1 receptors (Fig. 5E). Along with enhanced muscle regeneration, intrasoleus CDC-exosome injection decreased inflammation (Fig. 5F) and fibrosis (Fig. 5G). The net effect was complete restoration of contractile force in soleus muscles injected with CDC-exosomes (Fig. 5H).

**Figure 5.**
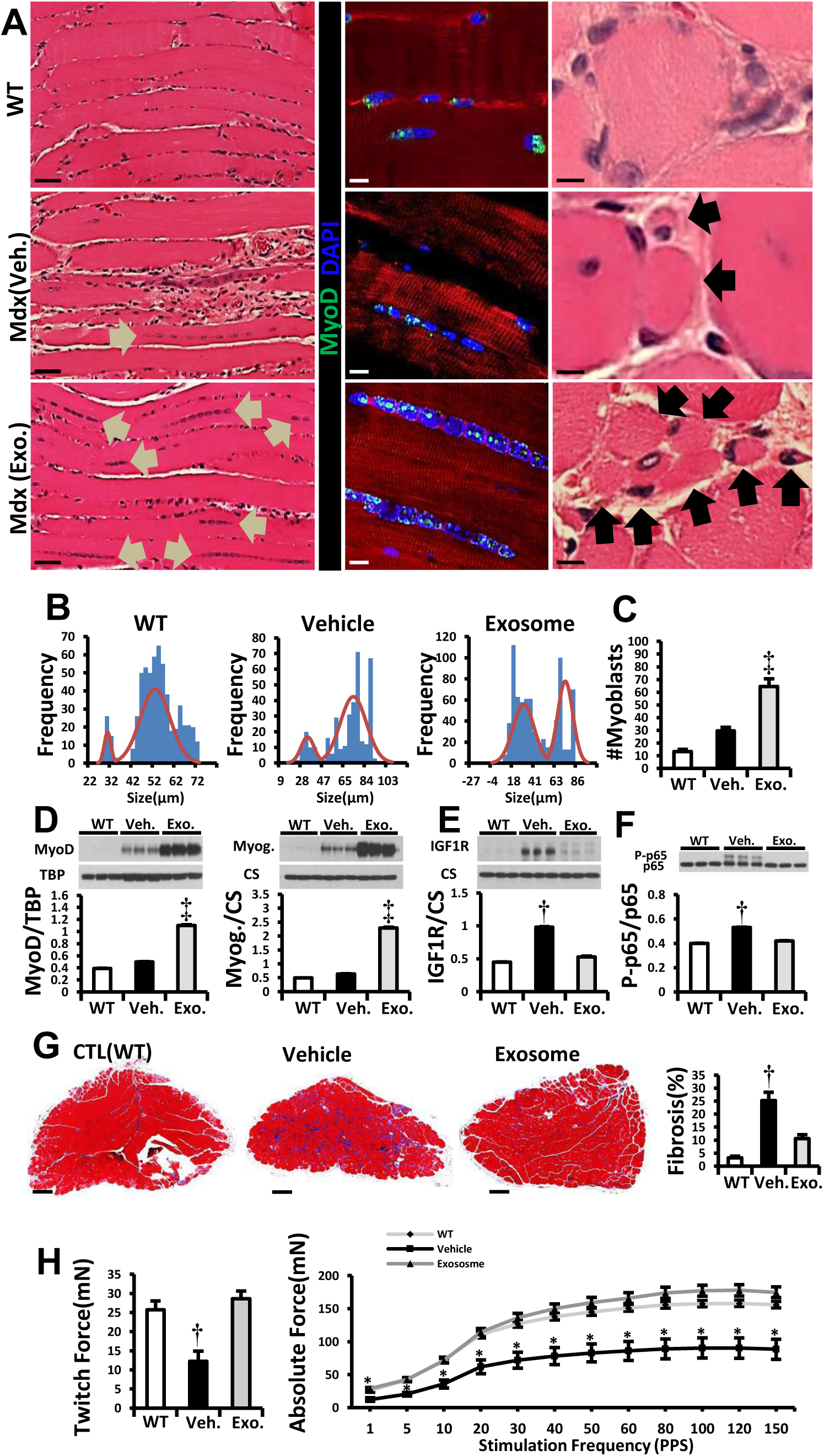
Intramuscular injection of CDC-exosomes resulted in muscle growth and reversal of pathophysiological abnormalities. A: H&E and immunohistochemical images of soleus muscle stained for MyoD (WT [wild type], vehicle and CDC-exosome-treated [*Mdx* (exosome)] *mdx* mouse soleus). Arrows in H&E images point to the lined up nuclei (left column) and myofibers (right column). In the immunohistochemistry, linearly-arranged nuclei were positive for MyoD (middle column). B&C: Frequency distribution of myofiber sizes and number of myoblasts (MyoD^+^) 3 weeks after vehicle/CDC-exosome injection in *mdx* soleus muscles (WT: n=5; Vehicle & Exosome: n=9 each). D-F: Western blots and pooled data for protein abundance of MyoD, myogenin (D), IGF1 receptor (IGF1R; E) and cytoplasmic p-p65 (F) in *mdx* soleus muscles 3 weeks after intrasoleus vehicle/CDC-exosome injection (WT: n=4; Vehicle & Exosome: n=6 each). G: Representative Masson trichrome images (enlarged in fig. S17) and morphometric analysis in *mdx* soleus muscles 3 weeks after administration of vehicle or CDC-exosomes into *mdx* soleus (WT: n=5; Vehicle & Exosome: n=9 each). H: *ex vivo* measurement of soleus contractile properties: twitch force and absolute force 3 weeks after vehicle/CDC-exosome injection into *mdx* soleus muscles. Pooled data are means ± SEM. *\*P< 0.05 vs. Mdx+CDC-exosome; †P< 0.05 vs. Mdx+CDC-exosome and WT (wild type mice); ‡P< 0.002 vs. Mdx+vehicle and WT (wild type mice). Scale bars: 5 μm (A, right column, 10μm (A, middle column), 50 μm (A, left column), 200 μm (G), 20 μm (H).*

### CDC-exosomes in human Duchenne cardiomyocytes derived from iPS cells

Demonstration of efficacy in multiple models of DMD would bolster the notion that CDC-exosomes may be viable therapeutic candidates. Duchenne human induced pluripotent stem cell (iPS)-derived cardiomyocytes (DMD CMs) exhibit a number of phenotypic deficits characteristic of DMD, including decreased oxygen consumption rate (OCR) reminiscent of that observed in *mdx* heart mitochondria (Fig. 1N), and abnormal calcium cycling (31). Priming DMD CMs with CDC-exosomes one week earlier suppressed beat-to-beat calcium transient alternans during 1Hz burst pacing (a measure of arrhythmogenicity (32); Fig. 6A) and normalized OCR (Fig. 6B). The congruence of experimental findings in the two DMD models is noteworthy: the *mdx* mouse has a nonsense mutation in exon 23 of the murine dystrophin gene, while the DMD patient whose iPS cells were studied here has a fundamentally different genetic lesion in the human dystrophin gene (exon 50 deletion with frame shift (31)). Thus, CDC-exosomes exert salutary effects in at least two classes of DMD mutations.

**Figure 6.**
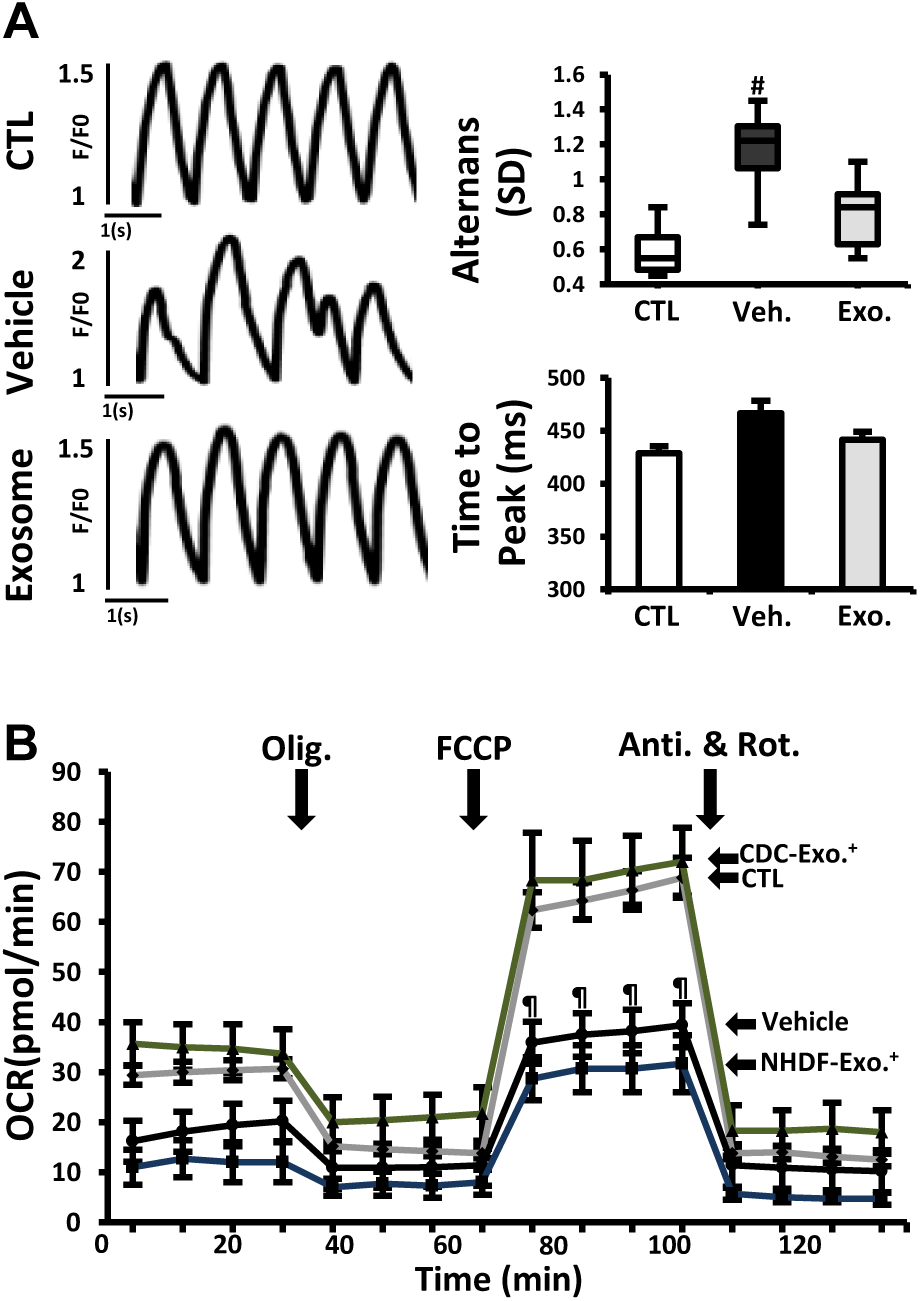
CDC-exosomes in human Duchenne cardiomyocytes derived from iPS cells. A: Calcium transients from normal and DMD CM measured during 1Hz burst pacing. Duchenne cardiomyocytes were primed with vehicle or CDC exosomes (exosomes) 1 week before assessment. Bar graphs of calcium transient alternans (variation in beat-to-beat calcium transient amplitude) and time to peak. B: Oxygen consumption rate (OCR) in DMD CM primed with CDC-exosomes or exosomes from normal human dermal fibroblasts (NHDF, as control; NHDF-exosome) 1 week before OCR measurement. Normal (CTL) and non-treated DMD CM (vehicle) were studied in parallel. See Fig. 1 legend for abbreviations. All data are means ± SEM except for the box plot (means ± SD). *\*P< 0.002 vs. CDC-exosome and CTL (Normal cardiomyocyte); #P< 0.03 vs. CDC-exosome and CTL (Normal cardiomyocyte); ¶ P< 0.02 vs. CDC-exosome.*

## DISCUSSION

We propose that CDCs act by secreting exosomes which are taken up by surrounding myocardium and by skeletal muscle, antagonizing multiple pathophysiological pathways that underlie DMD.

The congruent effects in *mdx* mice and in exon 50-deleted DMD hCM (Fig. 6) highlight the ability of CDC-exosomes to benefit multiple disease-causing dystrophin mutations. In the present work, we have not excluded the possibility that CDCs and their exosomes might, unexpectedly, increase dystrophin expression. We are presently investigating that possibility. Nevertheless, the dystrophin-independent actions of CDCs and their exosomal contents should be generalizable. After all, CDCs and their exosomes work quite well in cardiomyopathies not associated with dystrophin deficiency (7, 14, 27). The protean actions of CDCs and their exosomes include improved mitochondrial function, enhanced myocyte proliferation, and suppression of oxidative stress, inflammation and fibrosis (Fig. 7). Observed effects which appear to be independent of dystrophin restoration include the suppression of inflammation in *mdx* liver (an organ with no dystrophin expression). Notably, CDCs and their exosomes not only reverse the dystrophic phenotype but also forestall progression: the functional benefits of cardiac injections of CDCs or their exosomes persist for at least 3 months, and single doses of CDCs decrease mortality more than one year later in *mdx* mice. These findings beg the investigation of exosomally-triggered epigenomic modifications as a potential basis for the durable benefits (33), but such experiments are beyond the scope of this report.

**Figure 7.**
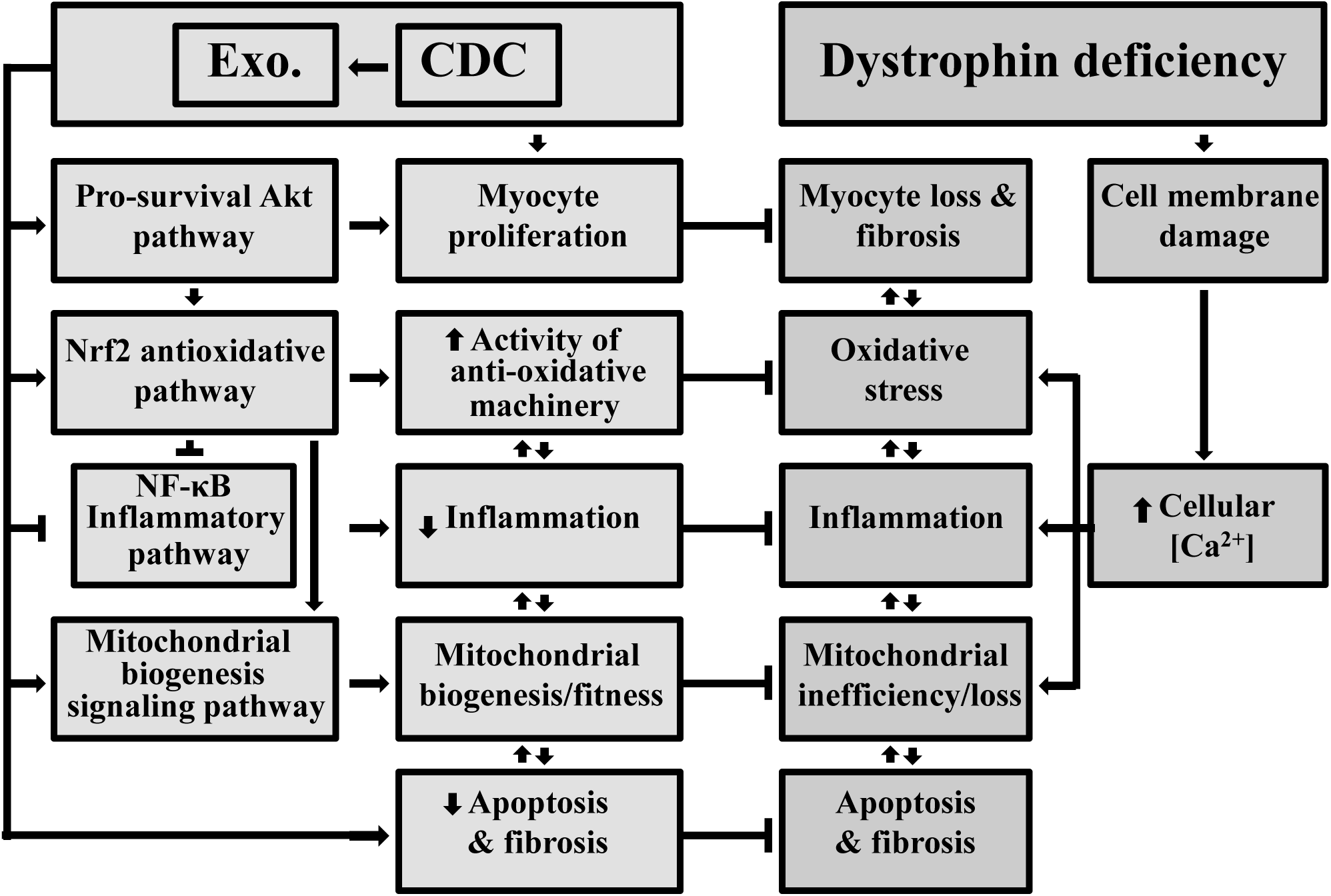
Schematic of pathophysiological mechanisms operative in Duchenne cardiomyopathy and the cellular mechanisms recruited by CDCs and their exosomes.

The ongoing HOPE-Duchenne trial is designed to assess single-dose delivery of CDCs to the heart (24). Whether in this first trial or in follow-up studies, our results give reason to wonder if CDCs will turn out to benefit not only the cardiomyopathy but also the skeletal myopathy of DMD. Exosomes derived from CDCs are promising next-generation therapeutic candidates with mechanisms of action fundamentally different to those currently in active development for DMD (e.g., exon skipping or myoediting).

## Methods

### Please see supplement for expanded methods

#### Animal study

We studied *mdx* mouse model of DMD (C57BL/10ScSn-Dmdmdx/J) and wild-type strain-matched mouse (C57BL/10ScSnJ wild type mouse heart) (Jackson Laboratory, USA) from 10 months of age. To optimize the process of CDC transplantation, preliminary dose-response experiments were performed, which identified 1x10^5^ cells in first injection and 1x10^4^ cells in second injection (3 months after first injection) as effective doses, consistent with prior dose-ranging experiments in ischemic and non-ischemic mouse models(14, 34). A total of 1x10^5^ cells/40μL phosphate-buffered saline (PBS; first injection) or 1x10^4^ cells/40μL PBS (second injection) or PBS alone were injected into left ventricular (LV) myocardium divided equally among 4 sites as described (14, 35). The LV was visually divided into three zones: basal, middle, and apical, with one injection in the basal, two in the middle and one in the apical zone. Ten-month-old CDC/*mdx* and vehicle/*mdx* mice were injected with CDCs (Mdx+CDC, n=12) or vehicle [placebo: Mdx+Vehicle (PBS), n=12] twice (3 months interval), respectively. Injections were during open-chest thoracotomy via a 28½ gauge-needle. All surgical procedures were carried out while the animals were under general anesthesia (Dexmedetomidine (0.5mg/kg)/Ketamine (75mg/kg); IP; once before surgery). Similar protocols were used for injection of CDC-exosomes into myocardium. Intraventricular single injection of CDC-exosomes [(10.32±3.28)x10^9^/150 μL PBS] or PBS alone into LV cavity were performed during open-chest thoracotomy via a 28½ gauge-needle. Intraaortic injections of CDCs (1x10^4^ cells/40μL PBS) or PBS were conducted using PE-10 catheter (ALZET; Cupertino, CA) via neck carotid artery. Intramuscular injection of exosomes into soleus (SOL) muscles were performed at a single site at the lower 1/3 of the muscle using a 25 μl Hamilton syringe (with 0.5 μl marks) with a 31 gauge needle. The needle was advanced up to the upper 1/3 of the muscle and then slowly retracted through the belly as exosomes [(20.64±2.12)x10^7^/3 μL] were injected. To ensure ethical and humane treatment of animals, we adhered to the principles recommended in the Guide for Care and Use of Laboratory Animals with oversight and approval by the Cedars-Sinai Health System Institutional Animal Care and Use Committee and the Department of Comparative Medicine (IACUC#3809).

### CDCs, CDC-exosomes, NHDF exosomes

#### CDC

Mouse CDCs were expanded from wild-type strain-matched mouse hearts (C57BL/10ScSnJ wild type mouse heart) as described(4). Briefly, ventricular tissues were minced into ∼1 mm explants, partially digested enzymatically and plated on adherent (fibronectin-coated) culture dishes. These explants spontaneously yield outgrowth cells (explant-derived cells) which were harvested once confluent and plated in suspension culture (10^5^ cells/mL on poly-D-lysine– coated dishes) to enable self-assembly of three-dimensional cardiospheres. Subsequent replating of cardiospheres on adherent culture dishes yielded CDCs which were used in all experiments at passage one.

#### CDC-exosomes

Exosomes were isolated from serum-free media conditioned overnight (24 hr) by cultured human CDCs(4, 7) (CDC-exosome) [or normal human dermal fibroblasts (NHDF) as a control] in hypoxia (2% O_2_; default condition) or normoxia (20% O_2_, solely for studies comparing RNA content of exosomes). Ultracentrifugation (100,000*g* for 1 hr) was used to isolate exosomes from conditioned media after sequential centrifugations at 300*g* (10min) and 10,000*g* (30min) and filtration with 0.22 micron filters(36). Isolated exosomes were re-suspended in PBS (for *in vivo* and *in vitro* experiments) and the ratio of exosome to protein was measured using Nanosight particle counter(37) and Micro BCA Protein Assay Kit (Life technologies, Grand Island, NY), respectively. Preliminary dose-response studies identified [(2.24±1.34)x10^7^] and [6.19±3.68 x10^8^] exosomes from hypoxic CDCs as effective doses for *in vitro* and *in vivo* (intramyocardial CDC-exosome injection) experiments, respectively. Exosomes were characterized by the most rigorous of criteria(38): linear iodixanol density gradient, transmission electron microscopy (TEM), key membrane proteins, and the biological effect. TEM images of sequentially-centrifuged exosomes with and without purification with linear iodixanol density gradient are shown (fig. S12A). The vesicles are variable in size and morphology, consistent with previous work(39). Western blot on lysed exosomes showed key proteins characteristic of exosomes: CD63, CD81 and TSG (fig. S12B). Biological activity of sequentially-centrifuged exosomes with (Exo1) and without (Exo2) purification with linear iodixanol density were compared by injection into *mdx* soleus muscles and evaluation of *mdx* soleus transcriptome 3 weeks after injection (fig. S12C). Correlation of fold changes in expression of same genes 3 weeks after Exo1 and Exo2 injection in *mdx* soleus muscles (fig. S12D) demonstrated similarity of the effects of Exo1 and Exo2 and supported the notion that the bioactivity of the vesicles isolated by our default protocol is genuinely due to exosomes, and not to another type of vesicles that might have been co-purified by ultracentrifugation.

### Echocardiography

Echocardiographic studies were performed two days before (Baseline) and 3 weeks, 2 and 3 months after first CDC/CDC-exosome (CDC-exosome) or vehicle injection and 3 weeks, 2 and 3 months after second CDC/CDC-exosome or vehicle injection (when applicable) using the Vevo 770 Imaging System (VisualSonics, Toronto, Canada)(4). The same imaging system was used to perform echocardiographic studies at baseline (2 days before) and 3 weeks after selected RNA (or control) injection. After induction of light general anesthesia, the heart was imaged at the level of the greatest LV diameter. LV ejection fraction (LVEF) was measured with VisualSonics version 1.3.8 software from 2-dimensional long-axis views.

### Treadmill exercise testing and survival analysis

For Fig. 1B, exercise capacity was assessed weekly with Exer-3/6 open treadmill (Columbus Instruments, Columbus, OH), beginning 1 week pre-operatively and 3 weeks after CDC/vehicle injection (exercise capacity measured in a subset of *mdx* mice 1 week pre-operatively was equivalent to that measured 3 weeks post-operatively in the Mdx+Vehicle group). After an acclimation period (10 m/min for 20 min), stepwise increases in average speed (2 m/min) were applied every two minutes during treadmill exercise until the mouse became exhausted (spending > 10 seconds on shocker; continuous nudging was used during treadmill to help mice stay on the track). Subsequently, the mouse was returned to the cage and the total distance recorded. The treadmill protocol conformed to guidelines from the American Physiological Society(40). After 3 months of weekly exercise, CDC/vehicle *mdx* mice along with wild-type age-matched mice were followed for assessment of mortality (Fig. 1C).

### In vitro isometric contractile properties of skeletal muscle

Mice were deeply anesthetized with Ketamine/Xylazine (80 mg/kg and 10 mg/kg body weight IP), the soleus (SOL) and/or extensor digitorum longus (EDL) and/or diaphragm (DIA) muscles were rapidly excised, and the animal was euthanized. Briefly, following a lateral midline skin incision of the lower leg the SOL and/or EDL muscle was dissected and isolated and its tendons of origin and insertion were tightened with silk suture (3-0) and rapidly excised. The SOL or EDL muscle was vertically mounted in a tissue bath containing a mammalian Ringer’s solution of the following composition: (in mM) 137 NaCl, 5 KCl, 2 CaCl_2_, 1 MgSO_4_, 1 NaH_2_PO_4_, 24 NaHCO_3_, 11 glucose.

The solution was constantly aerated with 95% O_2_ and 5% CO_2_ with pH maintained at 7.35 and temperature kept at 24°C. For studies of the diaphragm, following a left costal margin skin and muscle incision, a section of the midcostal hemidiaphragm was transferred to a preparatory Sylgar-lined dish containing cold Ringer’s and a narrow 3-4 mm wide strip of diaphragm was isolated maintaining fiber attachments to the rib and central tendon intact which were tighten with silk suture and mounted vertically in the tissue bath. One end of the SOL, EDL or DIA was secured to a clamp at the bottom of the dish and one end was attached to a calibrated force transducer (Cambridge Technology Model 300B, Watertown, MA). A micromanipulator linked to the system was used to adjust muscle length. Platinum plate electrodes placed on each side of the muscle were used for direct muscle stimulation (Grass Model S88 stimulator; Quincy, MA) using 0.2 msec duration monophasic rectangular pulses of constant current delivered at supramaximal intensity. Muscle length was adjusted until maximum isometric twitch force responses were obtained. Isometric contractile properties were determined at optimal length (Lo). Peak twitch force (Pt) was determined from a series of single pulses. Force/frequency relationships were measured at stimulus frequencies ranging from 5-150 pulses per second (pps). The stimuli were presented in trains of 1 sec duration with an interval of at least 1 min intervening between each stimulus train. Muscle forces generated, including Pt and maximum tetanic force (Po), were normalized for the estimated physiological cross-sectional areas (CSA) of the muscle segment (CSA = muscle weight/1.056 × Lo; where 1.056 g/cm^3^ represents the density of muscle) and expressed in Newtons (N)/cm^2^. For the SOL and EDL muscle Lo was also normalized for muscle fiber length (0.71 and 0.44 of Lo, respectively) in estimating muscle specific force. Absolute muscle forces generated by the SOL and EDL are also reported (mN)(41).

### iPSC derived cardiomyocytes

Urine-derived cells were seeded onto Matrigel (BD, San Jose, California) coated 12 well plates at 50,000 cells/well and allowed to attach overnight (day 0). On day two, cells were transduced with high-titer OSKM viral supernatants in the presence of 8 μg/ml polybrene for three hours. Viral supernatants were replaced with fresh USC medium and after three days, replaced with mTeSR1 medium (StemCell Technology, Vancouver, BC) and changed daily. As iPSC-like colonies appeared over time, they were picked using glass pasteur pipettes under a stereo dissection microscope (Leica M205C, Buffalo Grove, IL) and transferred to new Matrigel-coated plates for further expansion. Urine-derived iPSCs were differentiated to cardiomyocytes following an established protocol with modifications. Briefly, iPSC colonies were detached by 10 minute incubation with Versene (Life technologies, Carlsbad, CA), triturated to a single-cell suspension and seeded onto Matrigel-coated plastic dishes at a density of 250,000cells/cm^2^ in mTeSR1 medium and cultured for 4 more days. Differentiation was then initiated by switching the medium to RPMI-1640 medium supplemented with 2% insulin reduced B27 (Life Technologies) and fresh L-glutamine.

### Histology

Mice were sacrificed 3 weeks (WT: n=4; Mdx+Vehicle: n=6; Mdx+CDC/Mdx+CDC-exosome: n=6 each) or 3 months (WT: n=4; Mdx+Vehicle: n=6; Mdx+CDC/Mdx+CDC-exosome: n=6) after first CDC/CDC-exosome injections (n=6). Paraffin-embedded sections from apical, middle and basal parts of each heart or from diaphragm or entire soleus muscle (cross or longitudinal sections) were used for histology. Masson’s trichrome staining (HT15 Trichrome Stain [Masson] Kit; Sigma-Aldrich, St. Louis, MO) was performed for evaluation of fibrosis. T cells, B cells and macrophages were assessed by immunostaining with antibodies against mouse CD3, CD20 and CD68, respectively, and the average number of cells was calculated by counting cells in 10 fields from each of 10 sections selected randomly from the apical (3 sections; 50μm interval), middle (4 sections; 50μm interval) and basal (3 sections; 50μm interval) regions of each heart or from diaphragm. The data were presented as number of cells/mm^2^ field. Actively-cycling (Ki67^+^) and proliferating (Aurora B^+^) cardiomyocytes were counted in the same manner, and the cycling and proliferating fractions were expressed as the number of Ki67^+^, Aurora B^+^ divided by the total number of cardiomyocytes per high-power field (HPF), respectively, as described(14). Measurements were averaged for each heart. <u>Immunofluorescence staining:</u> Heat-induced epitope retrieval in low or high pH buffer (DAKO, Carpinteria, CA) was followed by 2 hours permeabilization/blocking with Protein Block Solution (DAKO, Carpinteria, CA) containing 1% saponin (Sigma, St. Louis, MO; Protein Block Solution contained 3% saponin was applied for immunofluorescence staining of Ki67). Subsequently, primary antibodies in Protein Block Solution were applied overnight in 4 C° for immunofluorescence staining of 5-μm sections from apical, middle and basal parts of each heart or cross sections of soleus muscle. After 3x wash with PBS, each 10 minutes, Alexa Fluor secondary antibodies (Life Technologies, Grand Island, NY) were used for detection. Images were taken by a Leica TCS SP5 X confocal microscopy system. Immunofluorescence staining was conducted using antibodies against Ki-67 (SP6; RM-9106-S0; 1:50, Thermo Fisher Scientific, Fremont, CA), WGA (Wheat germ agglutinin; W32466, 1:200; Thermo Fisher Scientific, Fremont, CA), Nrf2 (C20; 1:50; Santa Cruz Biotechnology, Santa Cruz, CA), aurora B (611082; 1:250; BD Biosciences, San Jose, CA). <u>Immunoperoxidase staining:</u>

Immunohistochemical detection of CD3 (AC-0004A), CD20 (AC-0012A) and CD68 (790-2931) was performed on 5-μm sections using prediluted rabbit monoclonal antibodies from Ventana Medical System (Tuscon, AZ; CD68) and Cell Marque (Rocklin, CA; CD3, CD20). Staining was conducted on the Leica Bond-Max Ventana automated slide stainer (Chicago, IL) using onboard heat-induced epitope retrieval method in high pH ER2 buffer (Leica Biosystems, Buffalo Grove, IL). Staining was visualized using the Dako Envision^+^ rabbit detection System and Dako DAB (Carpinteria, CA). The slides were subsequently counterstained with mayer’s hematoxylin for 1 minute and coverslipped. <u>Electron microscopy:</u> Apical (1 cube), middle (3 cubes from right, middle and left subparts) and basal (3 cubes from right, middle and left subparts) parts of posterior wall from each heart (WT: n=3; Mdx+Vehicle: n=3; Mdx+CDC: n=3) were fixed by immersion of 1mm^3^ cubes in 2% glutaraldehyde, postfixed in osmium, and embedded in epon. Sections were cut at silver thickness, stained with uranyl acetate and lead citrate, and viewed with JEOL 1010 equipped with AMT digital camera system.

### Western blots

Western blot analysis was performed to compare abundance of target proteins contributing to Nrf2 signaling [Nrf2, phospho-Nrf2 (Nrf2-p^s40^) and Nrf2 downstream gene products: heme oxygenase-1 (HO-1), catalase, superoxide dismutase-2 (SOD-2), and catalytic subunit of glutamate-cysteine ligase (GCLC)], Nrf2 phosphorylation [phospho-Akt(Akt-p^308^; Akt-p^473^)], oxidative phosphorylation [CI (NDUFB8 subunit), CII (SDHB subunit), CIV (MTCO1 subunit), CIII (UQCRC2 subunit) and CV (ATPSA subunit)], inflammation (NF-κB and MCP-1), fibrosis (Collagen IA1 and collagen IIIA1) and myofiber proliferation and differentiation (Myogenin, MyoD and IGF1R). Density of malondialdehyde protein adducts, a marker of oxidative stress, was also measured by Western blotting (WB). Samples from apical, middle and basal parts of each heart (each 1 mm-thick transverse section) were mixed and homogenized, and nuclear and cytoplasmic fractions were extracted per manufacturer’s instructions (CelLytic NuCLEAR Extraction Kit, Sigma-Aldrich, St. Louis, MO). The same kit was applied for extraction of nuclear and cytoplasmic fractions from entire soleus muscle, a narrow 3-4 mm wide strip of the left midcostal hemidiaphragm and iPSC derived cardiomyocytes. Mitochondria were extracted from fresh whole hearts (WT: n=3; Mdx+Vehicle: n=8; Mdx+CDC: n=8) as described in respirometry section. Cytoplasmic, nuclear and mitochondrial extracts for WB analysis were stored at −80C°. The protein concentrations in extracts were determined by the Micro BCA Protein Assay Kit (Life technologies, Grand Island, NY). Target proteins in the cytoplasmic, nuclear and mitochondrial fractions were measured by Western blot analysis using the following antibodies: antibodies against mouse Nrf2 (C-20; sc-722), HO-1 (H-105; sc-10789), catalase (H-300; sc-50508), SOD-2 (FL-222; sc-30080), GCLC (H-338; sc-22755), collagen IA1 (D-13; sc-25974), and collagen IIIA1 (S-17; sc-8780-R) were purchased from Santa Cruz Biotechnology (Santa Cruz, CA), phospho-Nrf2 (Nrf2-p^s40^; orb34864; Biorbyt, San Francisco, CA), respiratory chain subunits (Total OXPHOS Rodent WB Antibody Cocktail antibody; MS604), malondialdehyde (ab27642), citrate synthase (ab96600) and TBP (TATA binding protein; ab63766) [Abcam, Cambridge, MA], Akt (9272) and Akt-p^T308^ (5106S), IκB-α (4814S), p-IκB-α (9246S), phospho-NF-κB p65 (Ser536; 3033S) [Cell Signaling Technology, Denver, CO], MCP-1 (HPA019163), NF-κB p65 (SAB4502615) [Sigma-Aldrich, St. Louis, MO], Myogenin (F12B; MA5-11658), MyoD (5.8A; MA1-41017), Akt-p^S47^ (14-6; OMA1-03061) and IGF-IR/IGF1 Receptor (194Q13; AHO1292) [Thermo Fischer Scientific, Fremont, CA] antibodies were purchased from the cited sources. Antibodies to TBP and citrate synthase were used for measurements of the housekeeping proteins for nuclear (TBP), cytosolic and mitochondrial (citrate synthase) target proteins. <u>Western blot methods:</u> Briefly, aliquots containing 20 μg proteins were fractionated on 8, 10 and 4-12% Bis-Tris gel (Life technologies, Grand Island, NY) at 120 V for 2 h and transferred to a PVDF membrane (Life technologies, Grand Island, NY). The membrane was incubated for 1 h in blocking buffer (1× TBS, 0.05% Tween-20 and 5% nonfat milk) and then overnight in the same buffer containing the given antibodies at optimal dilutions listed in Suppl. Table 1. The membrane was washed 3 times for 5 min in 1× TBS, 0.05% Tween-20 before a 2-h incubation in a buffer (1× TBS, 0.05% Tween-20 and 3% nonfat milk) containing horseradish peroxidase-linked anti-rabbit IgG (7074P2), anti-mouse IgG (7076S) [Cell Signaling Technology, Denver, CO] and anti-goat IgG (A5420; Sigma-Aldrich, St. Louis, MO) at 1:1000-3000 dilution. The membrane was washed 3 times for 5 min in 1 × TBS, 0.05% Tween-20 and developed by autoluminography using the ECL chemiluminescent agents (Super Signal West Pico Chemiluminescent Substrate; Life Technologies, Grand Island, NY). Citrate synthase and TBP were used as housekeeping proteins against which expressions of the proteins of interest were normalized. Phosphorylated Akt, Nrf2 and IκB-α were normalized to total Akt, Nrf2 and IκB-α. Western blot analyses of collagen I and collagen III were conducted under non-reducing, non-denaturing conditions.

### Statistical analysis

All pooled data are presented as mean±SEM, except results for alternans data (Fig. 6A) which are presented as mean ±SD. Normality and equality of variances of data sets were first tested using Kolmogorov-Smirnov and Levene’s tests, respectively. If both were confirmed, t-test or analysis of variance followed by Bonferroni’s post hoc test were used for determination of statistical significance; if either normality or equality of variances was not assured, nonparametric tests (Wilcoxon test or Kruskal-Wallis test followed by Dunn’s post-test) were applied (SPSS II, SPSS Inc., Chicago, Illinois). No preliminary data were available for a power analysis. Results from a pilot project allowed us to power subsequent studies. The study followed preclinical reporting standards, as described (42). Age-matched mice were randomly allocated to experimental groups using computer generated randomization schedules (https://www.randomizer.org). Conduct of experiments and analysis of results and outcomes were performed in a blinded manner (allocation concealment and blinded assessment of outcome). There was no post-hoc exclusion of mice or data after the analysis before unblinding.

#### Ejection Fraction Data

Preliminary data were collected from a pilot study of 5 animals per group measuring ejection fraction at baseline, and again 3 weeks after treatment with cells or vehicle control in *mdx* and corresponding wild-type mice (C57BL/10ScSnJ; Suppl. Table 2). The measured treatment effect was approximately 4 units, with a time effect of approximately 1 unit, with group standard deviations of 3.5 units. We anticipated larger differences between groups over later time points with possible increase in measured variance. Therefore, with <u>12 animals per treatment group in the each of the</u> *<u>mdx</u>* <u>groups, and 7 wild-type control animals</u>, the study had at least 80% power to detect a difference of <u>4.5 units</u> or greater in treatment effect and <u>1.4 units</u> or greater in time effect in a study design with 6 measurements per animal over time assuming a compound symmetry covariance structure, a correlation of 0.7 between measurements within animals over time, and a two-sided alpha of 0.05. (Power computed via PASS v. 11.0.).

#### Treadmill Data

Preliminary data were collected from a pilot study of 5 animals per group measuring treadmill distance (i.e., the distance ambulated before exhaustion, as described below) at baseline, and again 3 weeks after treatment with cells or vehicle control in *mdx* animals and corresponding wild-type animals (Suppl. Table 2). The measured treatment effect was approximately 150 meters, with limited differences observed over time in untreated groups. Group standard deviations were approximately 75 meters, with more variation observed after treatment. We anticipated larger differences between groups over later time points with possible increase in measured variance. Therefore, with <u>11 animals per treatment group in the each of the transgenic groups, and 7 wild-type control animals</u>, the study had at least 80% power to detect a difference of <u>100 meters</u> or greater in treatment effect and changes of at least <u>30 meters</u> over time in a study design with 12 measurements per animal over time assuming a compound symmetry covariance structure, a correlation of 0.7 between measurements within animals over time, and a two-sided alpha of 0.05. (Power computed via PASS v. 11.0.).

## Acknowledgments

Supported by grants from Coalition Duchenne and NIH (R01 HL124074). We thank Liang Li for skilled technical assistance.

